# Phase variation in *Mycobacterium tuberculosis glpK* produces transiently heritable drug tolerance

**DOI:** 10.1101/717272

**Authors:** Hassan Safi, Pooja Gopal, Subramanya Lingaraju, Shuyi Ma, Carly Levine, Veronique Dartois, Michelle Yee, Liping Li, Landry Blanc, Hsin-Pin Ho Liang, Seema Husain, Mainul Hoque, Patricia Soteropoulos, Tige Rustad, David R. Sherman, Thomas Dick, David Alland

## Abstract

The length and complexity of tuberculosis (TB) therapy, as well as the propensity of *Mycobacterium tuberculosis* to develop drug resistance, are major barriers to global TB control efforts. *M. tuberculosis* is known to have the ability to enter into a drug-tolerant state, which may explain many of these impediments to TB treatment. We have identified a novel mechanism of genetically encoded but rapidly reversible drug-tolerance in *M. tuberculosis* caused by transient frameshift mutations in a homopolymeric tract (HT) of seven cytosines (7C) in the *glpK* gene. Inactivating frameshift mutations associated with the 7C HT in *glpK* produce small colonies that exhibit heritable multi-drug increases in minimal inhibitory concentrations and decreases in drug-dependent killing; however, reversion back to a fully drug-susceptible large-colony phenotype occurs rapidly through the introduction of additional insertions or deletions in the same *glpK* HT region. These reversible frameshift mutations in the 7C HT of *M. tuberculosis glpK* occur in clinical isolates, accumulate in *M. tuberculosis* infected mice with further accumulation during drug treatment, and exhibit a reversible transcriptional profile including induction of *dosR* and *sigH* and repression of *kstR* regulons, similar to that observed in other *in vitro* models of *M. tuberculosis* tolerance. These results suggest that GlpK phase variation may contribute to drug-tolerance, treatment failure and relapse in human TB. Drugs effective against phase-variant *M. tuberculosis* may hasten TB treatment and improve cure rates.

**SIGNIFICANCE:** The ability of *M. tuberculosis* to survive during prolonged treatment has been attributed to either transient stress responses or fixed heritable drug-resistance producing mutations. We show that phase-variation in the *M. tuberculosis glpK* gene represents a third type of resistance mechanism. The ability of these *glpK* mutants to grow slowly and then rapidly revert suggests that these transiently-heritable changes may also explain how a hidden population of drug-tolerant bacteria develops during TB treatment. As a genetically trackable cause of drug-tolerance, *M. tuberculosis glpK* mutants provides a unique opportunity to study these phenomena at a cellular and mechanistic level. These mutants could also be used for developing drugs that target tolerant populations, leading to more rapid and effective TB treatments.

## INTRODUCTION

Despite decades of control efforts, tuberculosis (TB) remains the leading cause of death from an infectious disease(1). The length and complexity of TB therapy is a major barrier to TB control. Drug-susceptible TB must be treated with multiple drugs, usually for 6 months, and multidrug-resistant TB must be treated for at least 9 months(2, 3). Relapses remain fairly common despite these regimens(4–7). Many of these clinical phenomena can likely be attributed to the ability of *Mycobacterium tuberculosis* to enter into a tolerant state when exposed to drugs, hypoxia, nutritional deprivation, or host defense mechanisms during human infections that renders them temporarily drug-resistant(8–10). This transient “phenotypic” drug-resistance is not thought to be caused by genetic changes associated with new heritable drug-resistance but has instead been attributed to reversible transcriptional and metabolic changes. A number of *in vitro* studies have confirmed that *M. tuberculosis* can become transiently drug resistant when cultured under growth-limiting conditions, including drug-treatment(11–13). Evidence for the development of reversible phenotypic drug-resistance during human TB treatment includes the observation that most patients who relapse after treatment for drug-susceptible TB remain infected with the same fully drug-susceptible strain(14–16). *M. tuberculosis* that has been cultured from apparently latent, closed and encapsulated lesions of patients undergoing TB treatment has also been found to be drug-susceptible(17).

Phase variation is an adaptive mechanism that mediates reversible switching “on” and “off” of a gene by genotypic changes such as DNA methylation, homologous recombination, DNA rearrangement, or insertions/deletions in short sequence repeats or homopolymeric tracts (HTs) located within the promoter region or the coding sequence of a gene(18). Reversible frameshift mutations in HTs are a result of slipped-strand mispairing errors during replication and are well documented in many bacterial species(19), with a high rate of HT variants observed in species with DNA mismatch repair (MMR) deficiency(20). *M. tuberculosis* also lacks a recognizable MMR system(21, 22). This suggests that the poly-G:C and poly-A:T tracts identified in the *M. tuberculosis* genome(21) may be susceptible to reversible insertion and deletion mutations during replication.

Here, we describe a novel mechanism of drug-tolerance that is caused by genetically encoded but rapidly reversible mutations in the 7C HT of the *glpK* gene in *M. tuberculosis*. These mutations produce small colony and morphological variants that have reduced susceptibility to drugs, but unlike classical drug-tolerance or persistence, are expressed as a transiently heritable trait. We propose that GlpK phase variability may account for much of the clinical and microbiological observations associated with persistence and relapse TB in humans.

## Results

### Small colonies with altered morphology are detectable in clinical *M. tuberculosis* strains

We examined cultures of 26 drug susceptible(23) and 8 drug resistant (1 Isoniazid (INH) and ethambutol (EMB) resistant, 1 Rifampicin (RIF) and INH resistant, and 6 RIF-INH-EMB resistant)(24) clinical *M. tuberculosis* strains. Of these, we noted that 9 susceptible and 4 RIF-INH-EMB resistant isolates contained a subpopulation of small colonies (SCs) with a smooth morphology mixed in with the large colonies (LCs) that predominated in the culture. We selected LCs and SCs of one pan-susceptible clinical isolate (C71) (Fig. 1*A*) and one multi-drug resistant clinical isolate (TDR193) (Fig. 1*B*). Whole genome sequencing (WGS) of the LC (C71-LC) and SC (C71-SC) isolates showed that they differed only at the *M. tuberculosis glpK* gene (*Rv3696c*, a glycerol kinase), after exclusion of the highly repetitive PE/PPE/PE-PGRS genes from the analysis. Compared to the *M. tuberculosis* H37Rv reference sequence (GenBank ID: AL123456.3)(21) (Table 1 and Fig. 1*C*), C71-LC had a 3C insertion in a 7C homopolymeric tract (HT) (*M. tuberculosis*, nucleotide 566 to 572) of *glpK*. In contrast, the C71-SC sequence contained both 3C and 4C insertions in the same 7C HT, indicating a sub-strain mixture within the colony. The 3C insertion resulted in a GlpK192Gly insertion that preserved the *glpK* open reading frame; however, the 4C insertion increased the size of the 7C HT to 11C and produced a *glpK* frameshift. We noted similar mutational differences between the LCs and SCs of the multidrug-resistant strain TDR193. WGS of the LC (TDR193-LC) and the SC (TDR193-SC) sequences of strain TDR193 only identified mutations in *glpK* and *rpoC* (Table 1). TDR193-LC had wild-type *glpK* and a nonsynonymous mutation in *rpoC*. Mutations in *rpoC* have been previously described as compensatory for the *rpoB* mutations associated with rifampicin resistance(25) and this mutation was not investigated further. TDR193-SC had a single C insertion in the *glpK* 7C HT causing a frameshift (Table 1 and Fig. 1*C*). We then sequenced this *glpK* hotspot in 94 additional *M. tuberculosis* clinical isolates randomly selected from the TDR-TB strain bank to represent a broad range of susceptibility profiles as well as geographic diversity(24) (*SI Appendix*, Table S1). The 7C HT was wild-type in 68, had a 1C insertion in 10, and showed a mixture of wild-type and 1C insertions in 16.

**Table 1.** Genotypic and phenotypic characteristics of large and small colonies of clinical *M. tuberculosis* strains.

| Strain | *embB* | *inhA* | *rpoB* | *rpoC* | *gyrA* | *gyrB* | *glpK* | Colony size |
| --- | --- | --- | --- | --- | --- | --- | --- | --- |
| TDR193-SC | Met306Val;<br>Met423Thr | Ile194Thr | Ser531Leu | Ala542Ala | Glu21Gln;<br>Ser95Thr;<br>Gly668Asp | Val340Leu | Ins573C<br>(7C→8C)<br>(frameshift) | Small |
| TDR193-LC | Met306Val;<br>Met423Thr | Ile194Thr | Ser531Leu | Ala542Ala;<br>Val775Ala | Glu21Gln;<br>Ser95Thr;<br>Gly668Asp | Val340Leu | WT (7C) | Large |
| TDR193-SC-RevLC1 | Met306Val;<br>Met423Thr | Ile194Thr | Ser531Leu | Ala542Ala | Glu21Gln;<br>Ser95Thr;<br>Gly668Asp | Val340Leu | Ins573CCC<br>(7C→10C)<br>(Gly192 insertion) | Large |
| TDR193-SC-RevLC9 | Met306Val;<br>Met423Thr | Ile194Thr | Ser531Leu | Ala542Ala | Glu21Gln;<br>Ser95Thr;<br>Gly668Asp | Val340Leu | WT (7C) | Large |
| C71-SC | Ala630Val | WT | WT | WT | Glu21Gln | WT | Ins573CCCC<br>(66%)/<br>Ins573CCC<br>(34%)<br>(7C→11C/10C)<br>(Mix of frameshift and Gly192 insertion) | Small and Large |
| C71-LC | Ala630Val | WT | WT | WT | Glu21Gln | WT | Ins573CCC<br>(7C→10C)<br>(Gly192<br>insertion) | Large |
| C71-SC-RevLC | Ala630Val | WT | WT | WT | Glu21Gln | WT | Ins573CCC<br>(7C→10C)<br>(Gly192<br>insertion) | Large |

**Fig. 1.**
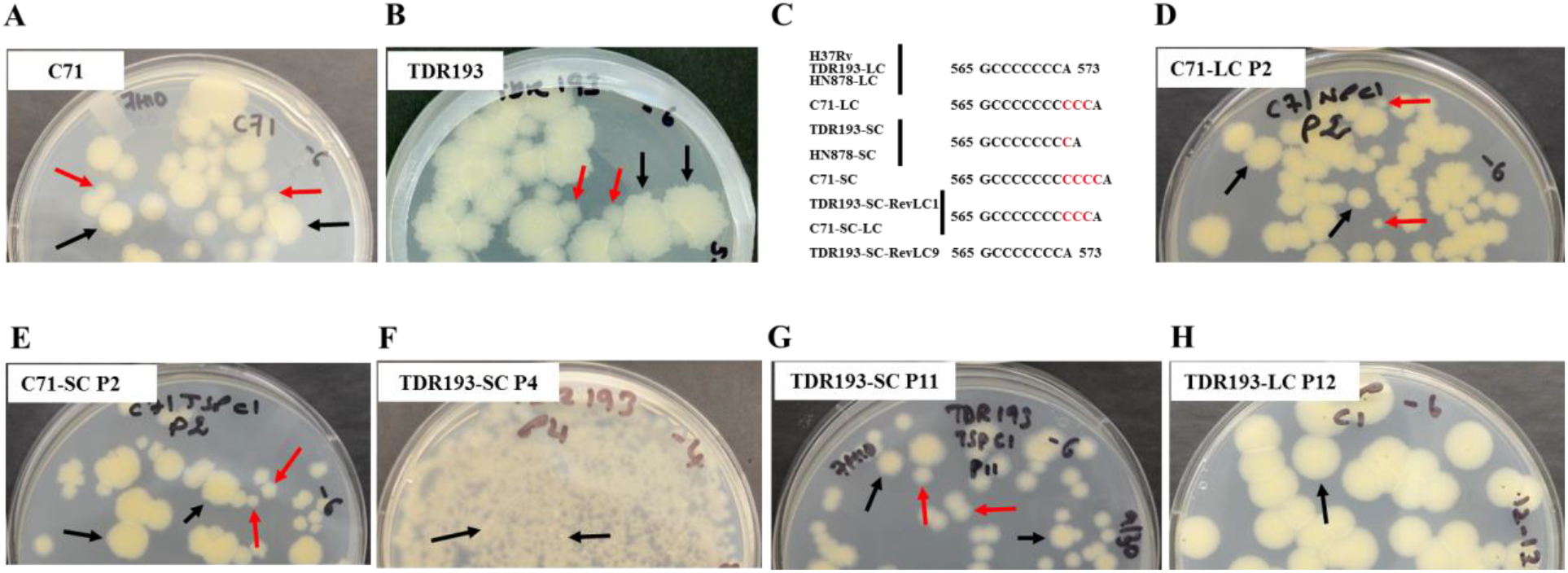
Clinical *M. tuberculosis* isolates with mixtures of small smooth and large rough colonies. Initial cultures of **(A)** Pansusceptible *M. tuberculosis* strain C71 and **(B)** multidrug resistant *M. tuberculosis* strain TDR193 were plated on 7H10 agar containing glycerol. (**C**) The *glpK* slippage site sequences of large and small colonies are shown. (**D**) C71-LC showed emergence of small colonies at passage 2. (**E**) C71-SC showed emergence of large colonies at passage 2. (**F**) Emergence at passage 4 and accumulation of large colonies at (**G**) passage 11 of a TDR193-SC culture. (**H**) TDR193-LC phenotype was stable after 12 passages. Examples of small colonies are indicated by red arrows and large colonies by black arrows.

### High frequency reversions among clinical *glpK* mutants

To determine whether clinical *glpK* SC frameshift mutants revert to normal colony size, we cultured TDR193-SC to stationary phase and plated the culture on glycerol containing agar to identify LCs. TDR193-SC cultures reverted to LC morphology at high frequency both in the presence or absence of glycerol in liquid medium (2.9 10^−2^±3.0 10^−2^ in 7H9 without glycerol vs 1.7 10^−2^±1.1 10^−2^ in 7H9 with glycerol). We also performed serial passage of C71-SC, C71-LC, TDR193-SC and TDR193-LC single colonies in the absence of glycerol. Both C71-LC and C71-SC showed a mixture of small and large colonies in just 2 passages (Fig. 1*D* and 1*E*), perhaps because of the instability of the large 10C and 11C *glpK* HT regions in these strains. In contrast, for TDR193, the SC (TDR193-SC) reverted to LC in 4 passages (Fig. 1*F-G*) but the LC (TDR 193-LC) was stable for 12 passages (Fig. 1*H*). WGS of one C71-SC colony that reverted to a LC phenotype (C71-SC-RevLC) revealed a deletion of 1C in the *glpK* HT, reducing the 11C HT to 10C. WGS of two TDR193-SC colonies that reverted to a LC phenotype revealed either a deletion of 1C in the *glpK* HT (TDR193-SC-RevLC9), reducing the 8C HT back to 7C as exists in the H37Rv reference strain, or the insertion of an additional 2C in the *glpK* HT (TDR193-SC-RevLC1) increasing the HT to 10C (Table 1 and Fig. 1*C*). Thus, each SC to LC mutant re-established the correct *glpK* gene open reading frame. No other new mutations were detected in the genomes of the revertants. Due to the high frequency of C71-SC and C71-LC reversions, compared to TDR193 variants, we did not perform any further experiments with C71 variants. Given the role of GlpK in glycerol metabolism, we cultured TDR193-SC, TDR193-LC, and TDR193-SC-RevLCs in media with or without glycerol supplementation. All strains had similar growth rates in liquid media (*SI Appendix*, Fig. S1*A*) and equal colony sizes on solid media (*SI Appendix*, Fig. S1*B*) in the absence of glycerol. However, in the presence of glycerol, strains with a functional *glpK* (TDR193-LC, TDR193-SC-RevLC1, and TDR193-SC-RevLC9) had improved growth in liquid media (*SI Appendix*, Fig. S1*A*) and produced larger colonies on solid media than TDR193-SC (*SI Appendix*, Fig. S1*B*). These results demonstrate that clinical *M. tuberculosis* strains frequently generate *glpK* 7C HT frameshift-variants that have a slower growth capacity under certain culture conditions and have the ability to rapidly revert to higher growth patterns by re-establishing a correct open reading frame.

### *M. tuberculosis* acquires frameshift mutations in the 7C HT of *glpK* during murine infections

We studied *glpK* mutant generation *in vivo* by infecting C56BL/6 mice with a low dose of *M. tuberculosis* strain HN878 via aerosol. One group of mice were not treated with any anti-tuberculars, and a second group received 100 mg/kg of Moxifloxacin (MXF) to simulate a human dose equivalent for one week (Fig. 2*A*). SC cells appeared to spontaneously arise, or to be selected for during infection and treatment (Fig. 2*B*). SCs were undetectable in pre-infection cultures of HN878. However, by week 7 post infection, SCs comprised 9% (± 5%) of the CFU recovered from the lungs of untreated control mice and 62% (± 9%) of the CFU recovered from the lungs of MXF treated mice (*p*=0.002) (Fig. 2*C*). WGS of 1 SC (named HN878-SC) randomly picked from lung cultures of untreated mice and 3 LCs (HN878-XM4-LC1, HN878-XM4-LC2, and HN878-XM4-LC3) and 3 SCs (HN878-XM4-SC4, HN878-XM4-SC5, and HN878-XM4-SC6) randomly picked from lung cultures of MXF treated mice was performed. Compared to the parental HN878 strain, no mutations were detected in all 3 LCs genome sequences. In contrast, all 4 SCs had a 1C insertion in the 7C HT of *glpK* (Fig. 1*C*), except for one SC (HN878-XM4-SC4) that also had a ΔC44 frameshift mutation in *Rv0452* and a synonymous Gly157Gly mutation in *purM*. To confirm the presence of *glpK* frameshift mutations in other SCs, we sequenced the *glpK* 7C HT in another 27 randomly picked SCs (12 from untreated and 15 from the MXF treated group) and 20 LCs (11 from untreated and 9 from the MXF treated group). All of 20 LCs had a wild-type *glpK* sequence while all 27 SCs had the same 1C insertion in the *glpK* 7C HT, leading to a frameshift. These results strongly suggest that *glpK* frameshift mutants arise during infection and have an improved ability to persist *in vivo*, particularly during antibiotic treatment.

**Fig. 2.**
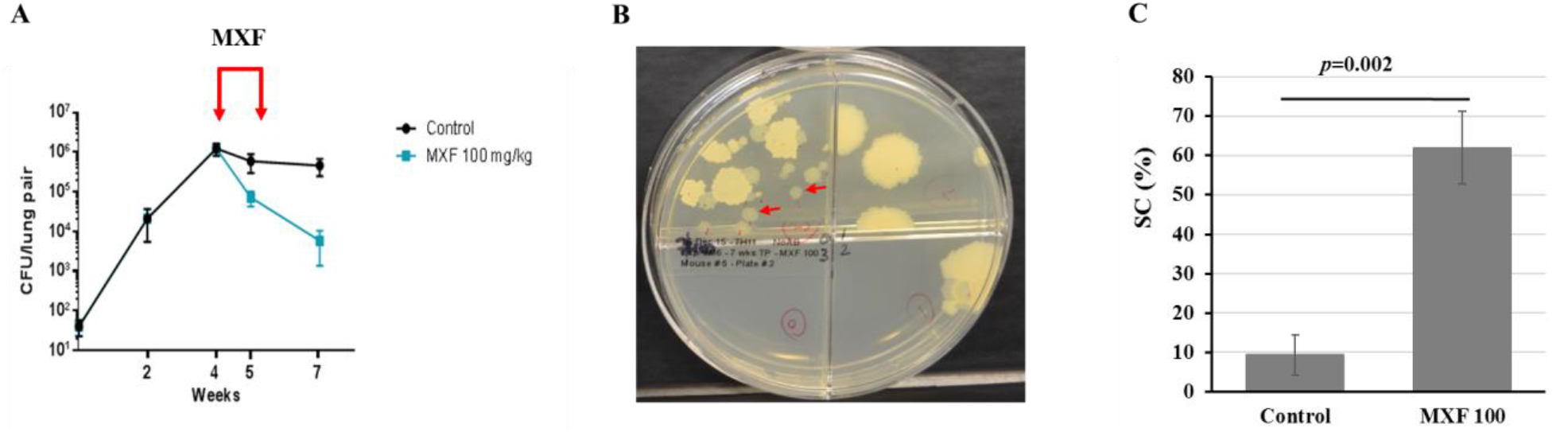
Frameshift *glpK* mutants develop in *M. tuberculosis* infected mice and are less susceptible to anti-tuberculosis treatment. **(A)** BALB/c mice were infected by aerosol with *M. tuberculosis* HN878 strain and treated with moxifloxacin (MXF 100mg/kg) over one week as indicated by red arrows. Shown are the averages and standard deviations of colony forming units (CFU) per lung of 4-5 mice sacrificed at the indicated time points. **(B)** Small smooth colonies isolated from an infected mouse lung after one week of MXF treatment. Examples of small colonies are indicated by red arrows. **(C)** Frequency, in percentage (%), of SC in the control and moxifloxacin treated groups of 4 mice each. Significant difference in frequency was determined by two tailed student’s *t* test.

### Small colony *glpK* frameshift mutants are less susceptible to sub-MIC concentrations of anti-tuberculosis drugs

Noting the accumulation of SCs during mouse infection and treatment, we tested SCs and LCs from various *M. tuberculosis* genetic backgrounds: clinical TDR193 isolate, HN878 identified after *in vivo* selection in mice, and SCs generated by performing clean *glpK* gene deletions in *M. tuberculosis* H37Rv, for alterations in drug susceptibility patterns. WGS analysis of the clinical multi-drug resistant TDR193-SC, TDR193-LC and TDR193-SC-RevLC9 strains identified no other mutations except as previously noted (Table 1). Despite these similarities, the INH MIC of TDR193-SC was 2-fold higher (5 μg/ml) than the MIC of TDR193-LC and TDR193-SC-RevLC9 (2.5 μg/ml) (Table 2). On the other hand, all variants had the same MICs for moxifloxacin (MXF), RIF, and EMB. However, the small colony TDRR193-SC had a 5 and 22-fold relative survival advantage measured by colony counts on sub-MIC concentrations of either MXF (0.1 μg/ml) or EMB (8 μg/ml), respectively, compared to LCs TDR193-SC-RevLC9 and TDR193-LC (Fig. 3*A*). The SC to LC revertant strain, TDR193-SC-RevLC9, had an intermediate survival advantage between the TDR193-LC and TDR193-SC. These differences at sub-MIC concentrations were only observed in the presence of glycerol. Next, we performed MIC experiments using a fully drug-susceptible HN878-SC strain isolated from our mouse study described above. An LC HN878 colony (HN878-LC) was used as a control after we had confirmed that this strain contained a wild type *glpK* gene. Both strains were isolated from the same lung of an infected mouse that had not been treated with MFX. These HN878 strains showed similar growth in media containing various concentrations of oleic acid (free fatty acid) or glyceryl trioleate (triglyceride derived from glycerol and three units of oleic acid, TAG), except in the presence of free glycerol where HN878-LC showed a more rapid growth rate (*SI Appendix*, Fig. S2*A*-*B*). HN878-SC and HN878-LC also had the same MICs for INH, RIF, EMB, and MXF, (Table 2). However, HN878-SC exhibited a 3 to 100-fold growth advantage, as measured by CFU counts, on sub-MIC concentrations of each of these drugs in the presence of glycerol, compared to HN878-LC (Fig. 3*B*). Finally, we created clean *glpK* knock out mutants in *M. tuberculosis* strain H37Rv (H37RvΔ*glpK*). The parental H37Rv and H37RvΔ*glpK* complemented with *M. tuberculosis glpK* inserted at the *M. tuberculosis attP* site and expressed under its native promoter served as controls. As with the HN878 SC strain, H37RvΔ*glpK* grew more slowly in liquid media supplemented with glycerol (*SI Appendix*, Fig. S2*C*), and produced smaller colonies on solid media supplemented with glycerol compared to H37Rv and the *glpK* complemented H37RvΔ*glpK* controls (*SI Appendix*, Fig. S2*D*). H37RvΔ*glpK* had identical INH, RIF and MXF MICs to both controls (Table 2). However, deletion of *glpK* in H37Rv enhanced strain survival in the presence of sub-MIC concentrations of all three antibiotics, similar to the studies performed with the *in vivo* selected HN878 SCs. This increased survival to anti-tuberculars was apparent both in the relative optical density of each strain cultured in various concentrations of INH, RIF and MXF compared to the same strain cultured in the absence of drug (Fig. 3*C-E*), and the relative survival of each strain plated on solid media containing 0.5 fold the MIC of INH, RIF and MXF compared to the same strains plated on solid media without drugs (Fig. 3*F*). The reversible SC phenotype that we observed in three different *M. tuberculosis* strains was not limited to the *M. tuberculosis* sensu stricto but was also observable in *M. bovis* BCG (BCG) (*SI Appendix*, Text S1 and Fig. S3).

**Fig. 3.**
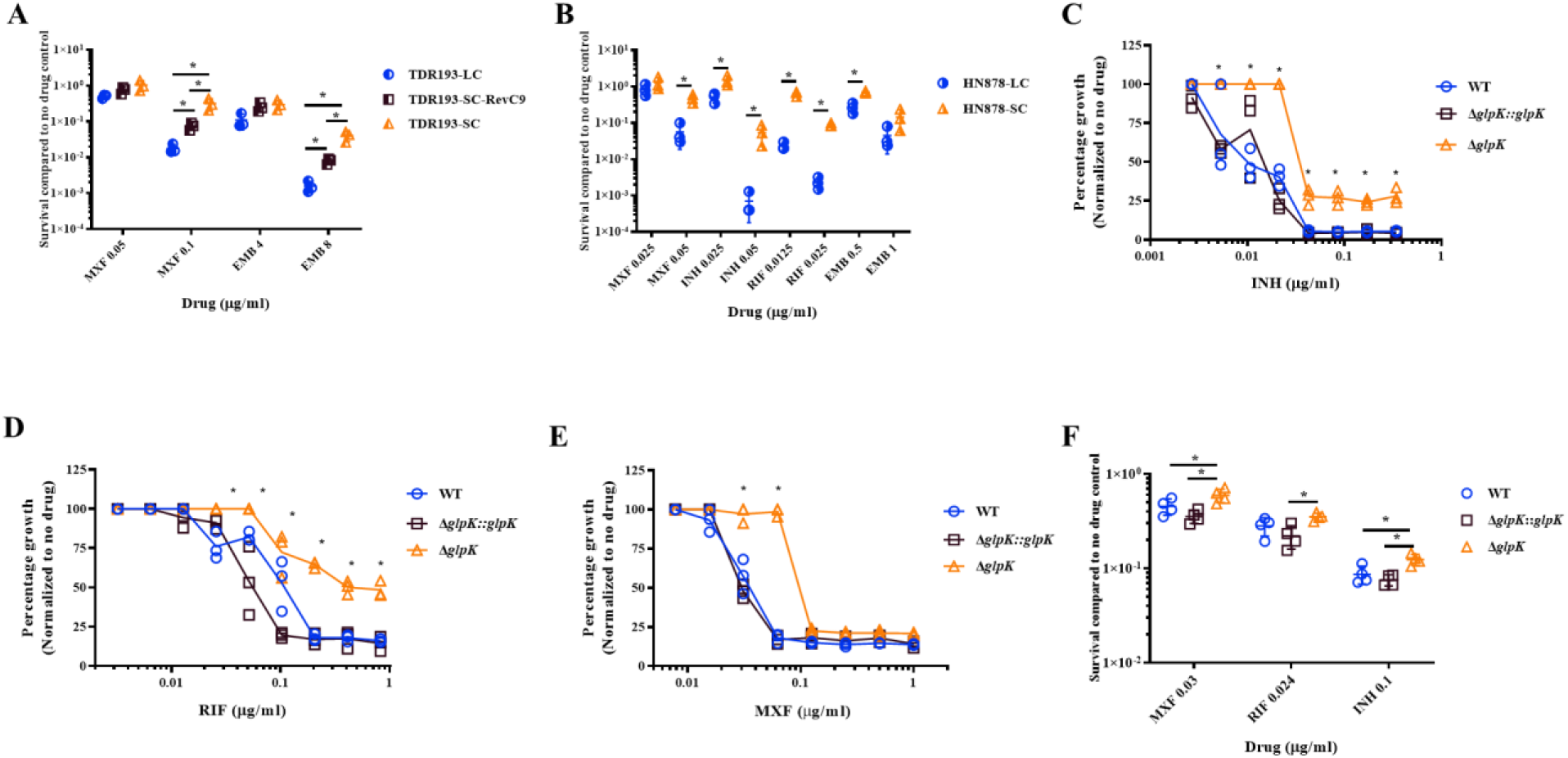
Decreased susceptibility of *glpK* mutants to various anti-tuberculosis drugs. Differential susceptibility of **(A)** TDR193 and **(B)** HN878 LC and SC variants to sub-MIC concentrations of MXF, EMB, INH and RIF were determined by percent survival of colony forming units (CFU) on 7H10 medium containing the antibiotic at the indicated concentrations versus no antibiotic control CFU. Differential susceptibilities of H37Rv wild type (WT), H37RvΔglpK and H37RvΔglpK∷glpK to sub-MIC concentrations of **(C)** INH, **(D)** RIF and **(E)** MXF compared to no antibiotic control were determined by OD_600_ measurements. **(F)** Susceptibility of H37Rv WT, H37RvΔglpK and H37RvΔglpK∷glpK to MXF, INH and RIF concentrations were determined by percent survival of CFU at the indicated concentrations compared to no antibiotic control. The values of three independent experiments are shown for each graph. Significant differences of survival frequencies were calculated using two tailed Student’s *t* test or two-way ANOVA analysis, \**p* <0.05.

**Table 2.** Minimum inhibitory concentration (MIC) of *M. tuberculosis* large and small colony variants.

| **Colony variant** | **MIC (µg/ml)** |  |  |  | **Resistance** |
| --- | --- | --- | --- | --- | --- |
|  | **MXF** | **INH** | **RIF** | **EMB** |  |
| **TDR193-LC** | **0.2** | **2.5** | **>100** | **16** | **MDR** |
| **TDR193-SC** | **0.2** | **5** | **>100** | **16** | **MDR** |
| **TDR193-SC-**<br>**RevLC9** | **0.2** | **2.5** | **>100** | **16** | **MDR** |
| **HN878-LC** | **0.1** | **0.1** | **0.05** | **2** | **Susceptible** |
| **HN878-SC** | **0.1** | **0.1** | **0.05** | **2** | **Susceptible** |
| **H37Rv** | **0.063** | **0.042** | **0.048** | **1.28** | **Susceptible** |
| **H37RvΔ*glpK*** | **0.063** | **0.042** | **0.048** | **1.28** | **Susceptible** |
| **H37RvΔ*glpK*::*glpK*** | **0.063** | **0.042** | **0.048** | **1.28** | **Susceptible** |

### Small colony *glpK* frameshift mutants are tolerant to supra-MIC concentrations of anti-tuberculosis drugs and hydrogen peroxide

We tested the extent to which SC *glpK* mutants exhibit a typical tolerant-like phenotype(26, 27). Performing time-kill studies, we found that the HN878-SC (*glpK* mutant) had delayed killing compared to HN878-LC cultures incubated with either 0.4μg/ml of MXF (4X the MIC) or 1μg/ml of INH (20X the MIC) (Fig. 4*A-C*). When incubated in MXF, the SC showed a 14-fold increase in survival on day 3 (*p*<0.01) and a 12-fold increase on day 5 (*p*<0.01). When incubated with INH, the SC showed a 2.84-fold increase in survival on day 1 (*p*<0.01) and a 2.72-fold increase on day 3 (*p*=0.05), compared to HN878-LC (*glpK* WT), in the presence of glycerol. We also tested the susceptibility of HN878-SC to the oxidative stress-inducing agent hydrogen peroxide. The HN878-SC was significantly more resistant to hydrogen peroxide (175mM), in the presence of glycerol, compared to HN878-LC (Fig. 4*D*).

**Fig. 4.**
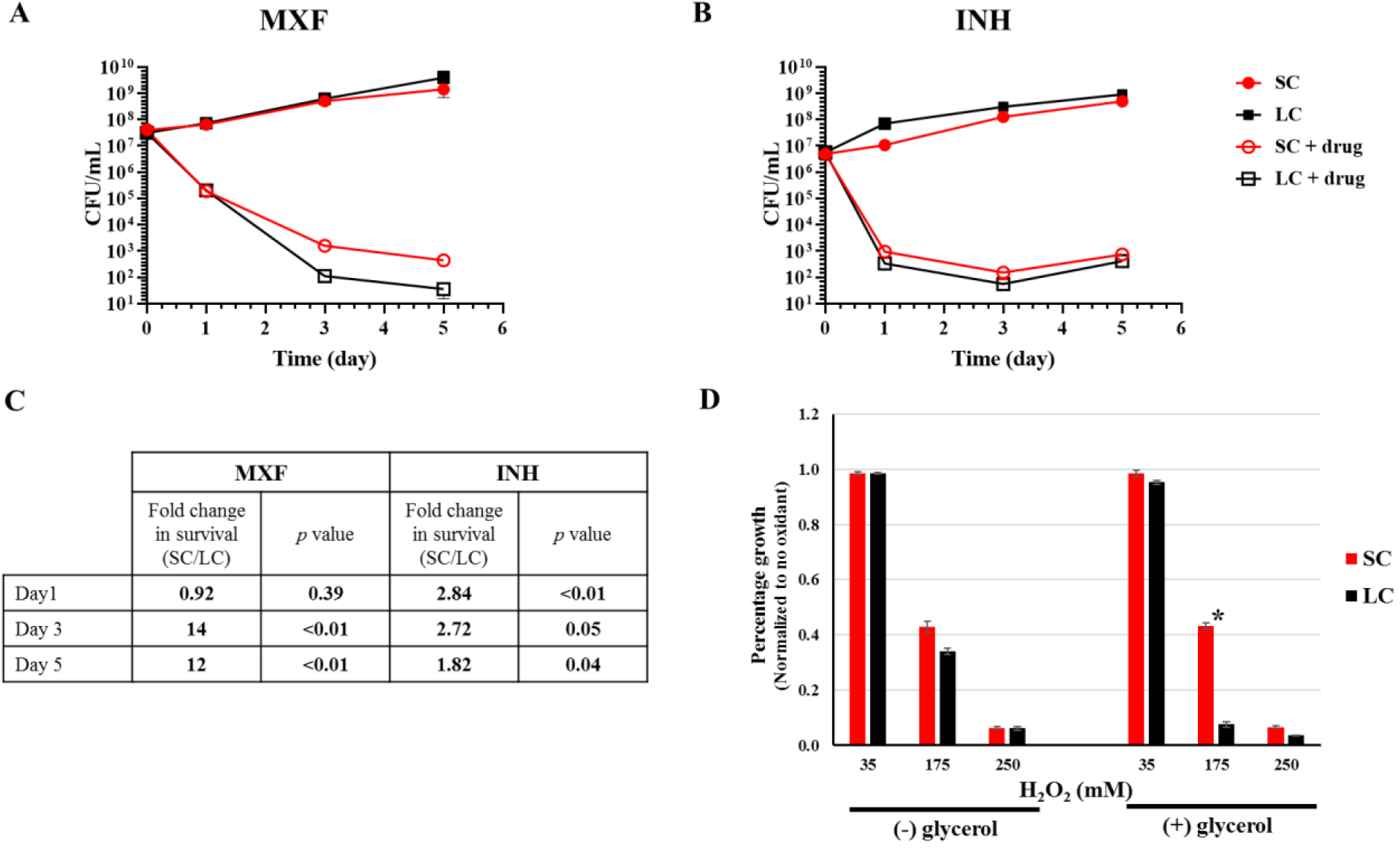
Differential tolerance of HN878 SC and LC variants to supra-MIC concentrations of antibiotics and to hydrogen peroxide. *M. tuberculosis* HN878 SC and LC strains grown in 7H9 supplemented with OADC and Tween 80 to mid-log phase (OD_600_ =0.6 to 0.7) were diluted to approximately 5 × 10^6^ or 5 ×10^7^ CFU/ml and treated in the presence of glycerol with **(A)** 0.4μg/ml MXF (4x MIC) or **(B)** 1μg/ml INH (20x MIC). Viability was determined by CFU counts. **(C)** Fold change with *p*-value of survival differences between SC and LC identified in panel A and B. **(D)** Differential susceptibility of HN878 SC and LC strains to H_2_O_2_ compared to control was determined by OD_600_ measurements. Both strains were grown in 7H9 medium containing OADC and Tween 80 with (+) or without (−) glycerol to mid-log phase (OD_600_ = 0.6 to 0.7). The cultures were diluted 1/20 and treated with different amounts of H_2_O_2_. The treated and control cultures were incubated for 6 days at 37°C. OD_600_ were recorded and normalized to the corresponding control without oxidant treatment. The values of three independent experiments are shown for each graph. Significant differences of survival frequencies were calculated using one tailed Student’s *t* test, * *p*<0.05.

### Transcriptional profiling of *M. tuberculosis glpK* 7C frameshift mutants

To identify altered transcriptomic pathways highly associated to *glpK* mutation, we performed RNA-Seq analysis of SC and LC variants with different genetic *M. tuberculosis* backgrounds, namely the SC strain TDR193-SC and the LC strain TDR193-SC-RevLC9 (lineage 4, Euro/American family), and the HN878-SC and HN878-LC strains (lineage 2, Beijing family), both in the absence and presence of glycerol. In both *M. tuberculosis* strain backgrounds, the SCs showed gene expression patterns similar to that of the LCs in the absence of glycerol (Fig. 5*A*). By contrast, the LCs cultured in the presence of glycerol differed by coordinated expression changes associated with responses typically associated with adaptation to diverse stressors. For example, several stress response regulons, including SigH(28) and DosR(29), were upregulated in the SCs with or without glycerol, relative to the LC strains in glycerol, while KstR(30) was among regulons downregulated in the SCs. Conversely, transcriptions of the genes involved in triacylglyceride synthesis were elevated in SC strains compared to their LC controls (Fig. 5*B* and *SI Appendix*, RNAseq *SI*). Collectively, these transcriptional changes are consistent with the possibility that *glpK* gene expression promotes *M. tuberculosis* growth under glycerol-replete conditions, whereas loss of functional *glpK* results in *M. tuberculosis* general stress responses associated with an adaptive tolerant state in the presence of glycerol.

**Fig. 5.**
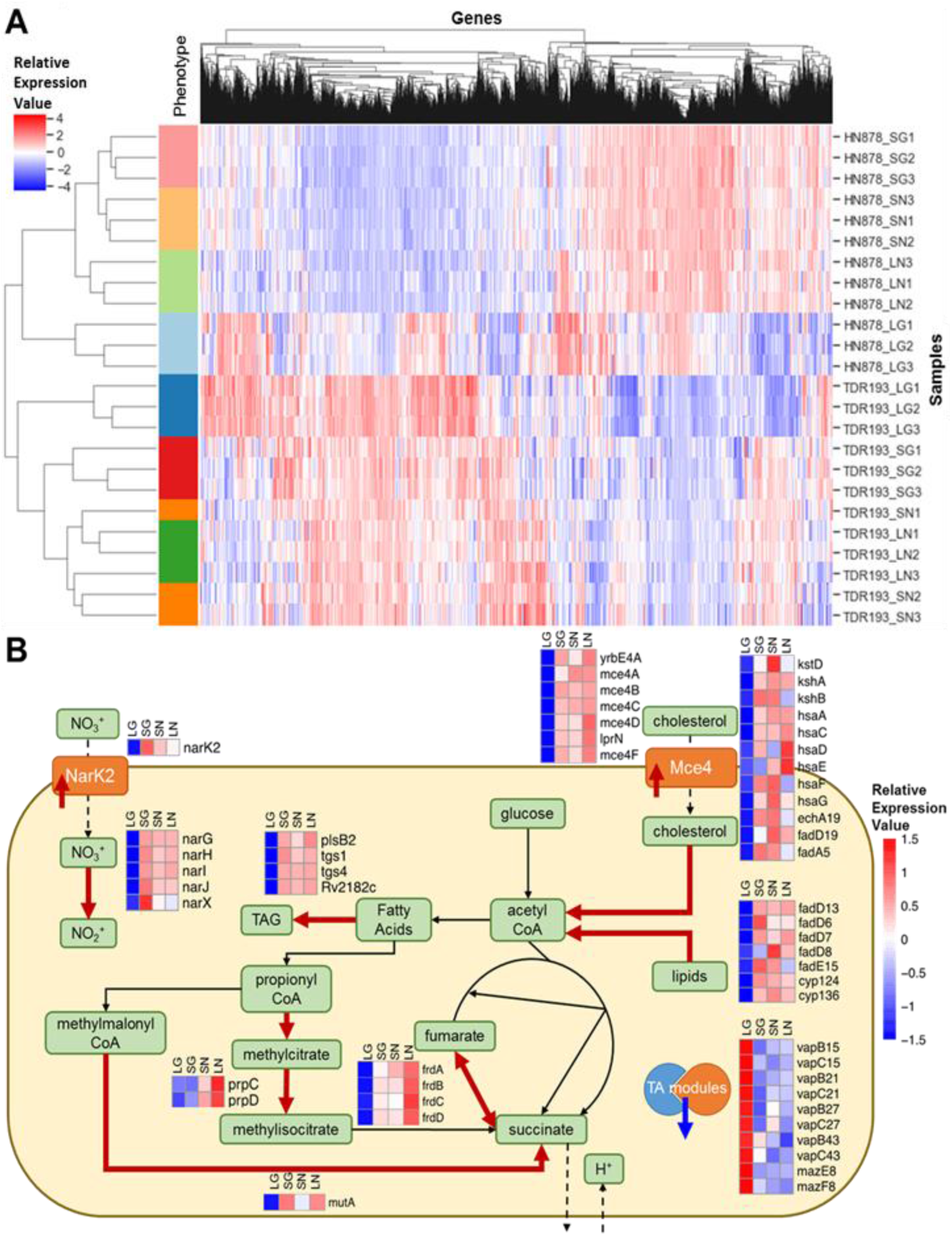
Gene expression differences appeared between the LCs and SCs in presence and absence of glycerol. **(A)** Heatmap of ranked gene expression values for the different conditions profiled by RNA-Seq. Individual samples are arrayed in the rows, and genes are arrayed along the columns. The different conditions profiled are: SG = small colony in glycerol (red bars); SN = small colony without glycerol (orange bars); LG = large colony in glycerol (blue bars); LN = large colony without glycerol (green bars). The HN878 strain background is represented in the light color bars, and the TDR193 strain background is represented in the dark color bars. Values reported are ranked RPKM values, scaled such that the mean across samples is 0 and the standard deviation is 1. The hierarchical clustering by sample shows clear separation by condition within each strain background. The largest separation between conditions is the LG vs. the other conditions. **(B)** Schematic summarizing main bacterial cellular processes impacted by glpK phase variation, as implied by gene expression changes in both TDR193 and HN878 backgrounds. The heatmaps visualize the scaled means of ranked RPKM values for the corresponding genes in each condition. Thick red arrows indicate gene associated proteins or processes that would be upregulated in SCs and in absence of glycerol, and the thick blue arrow indicates downregulated proteins.

### Lung glycerol in *M. tuberculosis* infected mice

To assess whether *M. tuberculosis* may encounter glycerol in human-like TB lesions, we tested the lungs of *M. tuberculosis* infected C3HeB/FeJ mice for the presence of glycerol using LC-MS. We used this infection model because C3HeB/FeJ mice develop much more human-like lung granulomas exhibiting caseous necrosis following aerosol infection with *M. tuberculosis*(31) than is observed in C57BL/6 mice(32). Glycerol was detectable in normally appearing lung tissue (61.6 μg/ml), lung tissue with cellular infiltrates (44.13 μg/ml) as well as in lung caseum (10.4 μg/ml), with ~10 to 30% of the glycerol extracted from triglycerides. These results suggest *in vivo M. tuberculosis* infections may be exposed to glycerol that may both stimulate increased growth in wild type *glpK* variants, and also select for *glpK* mutants where slower growth or drug-tolerance may improve survival.

## Discussion

Drug resistant *M. tuberculosis* has long been thought of as a stable phenotype that occurs through acquisition of heritable mutations. In contrast, drug-tolerance and drug-persistence have been viewed as reversible phenomena associated with transient phenotypic states. Here, we show in clinical strains of *M. tuberculosis* a novel type of drug-tolerance that is caused by a rapidly reversible *glpK* frameshift mutation in a 7C HT sequence, which produces a phenotype that is similar to the traditional definition of drug-tolerance(26, 27). These tolerance-conferring frameshifts also slow bacterial growth and produce small colonies. The *glpK* mutant small colonies showed small but significant increases in survival following exposure to various anti-tuberculosis agents, compared to *glpK* wild-type large colonies. Although these increases are small, such differences were consistent across different genetic *M. tuberculosis* backgrounds.

The association between genotypic drug-tolerance and slow growth, combined with a rapid reversal rate has potentially far reaching consequences, in that this process can mimic a drug-tolerant or persistent phenotype. For example, in the case of TB relapse the development of reversible drug-tolerance during TB treatment could permit a slow-growing bacterial subpopulation to survive drug treatment while causing minimal clinical symptoms. With the lifting of drug pressure at the end of treatment, the slow-growing population could rapidly revert to a wild-type population with the capacity to overgrow any drug-tolerant small colony cells, making the drug-tolerant subpopulation difficult to detect by conventional microbiological methods unless specifically assessed as in this study. Relapse in such a patient would thus be attributed to re-growth of a (drug-susceptible) “persister” population rather than genotypic drug-tolerant mycobacteria. Our findings are supported by a previous study that identified *glpK* frameshift mutations in a screen for mutations that increase tolerance to streptomycin and rifampicin in *M. tuberculosis*(13). Also demonstrating the potential clinical impact of even small changes in MIC, a recent study showed that increases as little as 0.01 μg/ml INH or RIF MIC in fully drug-susceptible *M. tuberculosis* isolates substantially raised the odds ratio for clinical relapse after TB treatment(33).

Our discovery that drug-tolerance is linked to slow growth and glycerol metabolism suggests an important role for nutritionally controlled differences in growth rates in TB pathogenesis. Earlier studies have shown that *M. tuberculosis* growth is strongly enhanced by adding glycerol to the culture medium compared to carbon sources such as glucose or free fatty acids. This has led to the use of glycerol in all standard mycobacterial media(34). Upon uptake, glycerol is phosphorylated by GlpK to glycerol-3-phosphate, which is either utilized for the synthesis of glycerophospholipids(35) or alternatively is converted by glycerol-3-phosphate dehydrogenase to dihydroxyacetone phosphate, which then enters glycolysis and gluconeogenesis(36). Furthermore, GlpK belongs to the ROK (repressor, open-reading frame, kinase) protein family(37); therefore, it is also possible that GlpK (as shown for other sugar kinases(38)), might also moonlight as a transcription regulator. When *M. tuberculosis* is starved for glycerol by either frameshifting *glpK* or by depleting glycerol from the medium, we suggest that a general stress response is activated, which phenotypically renders *M. tuberculosis* broadly stress resistant and antibiotic-tolerant. The RNA expression pattern that we noted to be part of this nutrient-limited stress response appears to be similar to that reported for wild-type *M. tuberculosis* cultured under conditions that have been associated with a “tolerant” “persistent” state(29). Culture of the LC but not the SC in rich glycerol media caused LCs to increase their growth rate. This increased growth and the accompanying down-regulation of stress regulons typified by DosR and SigH may have made the LC cultures more broadly susceptible to drugs and hydrogen peroxide. Therefore, *M. tuberculosis* appears to induce a general stress response by toggling on and off the expression of the *glpK* gene via reversible frameshifts within a 7C HT sequence.

We noted that all of the SC *glpK* mutants were caused by frameshift mutations occurring at a 7C HT located within the coding region of *glpK*. HTs in MMR-deficient bacteria are known to present slippage sites for DNA polymerases(39, 40), which produce frameshift mutations at high frequency(41, 42). *M. tuberculosis* also lacks a recognizable MMR system(21, 22). Furthermore, the *M. tuberculosis* H37Rv reference sequence (GenBank ID: AL123456.3)(21), includes 126 homopolymeric sequences (poly-C:G and poly-A:T ≥ 7-mer) located within open reading frames and 18 located within intergenic regions. A search of 5604 *M. tuberculosis* genomes in the NCBI database found at least one frameshift mutation in 74% of these homopolymeric sequences (*SI Appendix*, Table S2). However, this mutational mechanism to our knowledge has not been previously reported as a cause of reversible drug-tolerance. Previous studies of numerous pathogenic bacteria such as *Neisseria* species, *Mycoplasma* species, and *M. abscessus* have described the occurrence of frameshift mutations via slipped-strand mispairing in regions even without homopolymeric sequences(18, 43, 44). These findings suggest that reversible frameshift mutations as a means of genetically regulating stress responses, including antibiotic tolerance, may be widespread in multiple organisms including *M. tuberculosis*.

We showed that *glpK* frameshifted SCs develop during murine infections and that selection increases further with drug treatment. Our study was performed in mice receiving a single drug (MXF), because drug tolerant *M. tuberculosis* in human TB is likely to develop in diseased lesions where poor drug penetration often results in effective monotherapy(45). We found that *glpK* 7C HT variants emerged at a high rate under this circumstance. Pethe et al. showed that pyrimidine-imidazoles (PIs), whose mechanism of action is linked to glycerol metabolism do not inhibit *M. tuberculosis* infection in a murine model(46). However, the relative proportion of LC and SC generated in PI treated mice was not noted or recorded(46). We suspect that it is possible that PI treatment provided a strong selection for the emergence of *glpK* mutants in these mice. Variants with *glpK* frameshift mutations are detectable in sputum samples from many human TB patients and, furthermore, these *glpK* mutants were unstable(47, 48). Human plasma and mouse lung tissue contain significant amounts of free-glycerol(49–52) and our results show that free glycerol is also present in diseased mouse lung and caseum. The levels of glycerol we detected *in vivo* are substantially lower than those used in *M. tuberculosis* culture medium. While the level of glycerol needed to promote a growth difference between WT and *glpK* mutants *in vivo* is unknown, we have shown that the *in vivo* environment provides strong selection for *glpK* mutations. Overall, our results strongly suggest that SC emergence and drug-tolerance caused by GlpK phase variation is relevant in human TB. We also suggest that a larger repertoire of SCs caused by reversible frameshift mutations elsewhere in the *M. tuberculosis* genome may provide this pathogen with multiple biological mechanisms to adapt to drugs and changing environments.

## Methods

### Bacterial strains and culture conditions

Clinical *M. tuberculosis* strains were randomly selected from a collection of isolates obtained from the Tuberculosis Trials Consortium of the Centers for Disease Control and Prevention conducted Study 22(23) and from TDR-TB strain bank established by the World Health Organization Special Programme for Research and Training in Tropical Disease using geographic, phylogenetic and drug-resistance diversity as selection criteria(24). Unless otherwise stated, the *M. tuberculosis* strains were cultivated at 37°C either in Middlebrook 7H9 broth (Difco) containing 0.05% Tween 80 or on Middlebrook 7H10 agar supplemented with 0.5% glycerol, both enriched with 10% oleic acid-albumin-dextrose-catalase (Difco). Broth cultures were incubated under gentle shaking. *M. bovis* BCG (ATCC 35734) wild-type and mutant strains were maintained in complete Middlebrook 7H9 medium (BD Difco) supplemented with 0.05% (vol / vol) Tween 80, 0.5% (vol / vol) glycerol, 0.5% albumin, 0.2% glucose, 0.085% sodium chloride and 0.0003% catalase at 37°C with agitation at 80 rpm. Pyrazinoic acid (POA) was purchased from Sigma-Aldrich and was freshly dissolved in 90% DMSO at a concentration of 0.5 M and sterilized using 0.2 μm PTFE membrane filters (Acrodisc PALL). The POA-resistant *M. bovis* BCG strains used in this study were isolated previously as described in Gopal et al., 2016(53). For all plasmid construction, *Escherichia coli* strains Top10 (Invitrogen) were grown in Luria-Bertani broth or agar (both from Sigma Aldrich) at 37°C, supplemented with 50 μg/ml kanamycin (Sigma Aldrich) or 150 μg/ml Hygromycin B (Invitrogen), where appropriate.

### Deletion of *glpK* gene from *M. tuberculosis* H37Rv strain

The *glpK* gene was deleted using allelic exchange techniques as described previously(54). Briefly, 1500 base pairs upstream and downstream of *glpK* were cloned into p2NIL suicide vector containing *lacZ-sacB* selection cassette. All cloning were done in *E.coli* XL10 Gold (Aglient) and the final mutant construct was confirmed by Sanger sequencing. H37Rv strain was transformed with the final mutant construct followed by two-step selection process, as described previously(54). The *glpK* deletion was screened by PCR and confirmed by whole genome sequence as described below.

To complement the *glpK* deletion with the gene transcribed from its own promoter, a fragment containing the *glpK* gene and 200 bp upstream and 50 bp downstream sequences was cloned into pMV306 integrative plasmid. The resulting construct was electroporated into the *glpK* mutant and hydromycin resistant transformants were selected.

### Minimum Inhibitory Concentrations determination

*M. tuberculosis* strains were grown in 7H9 medium to mid-log phase (OD_580_=0.5-0.7). The cultures were then diluted to approximately 2 × 10^4^ CFU per starting inoculum and spotted on 7H10 agar plates containing the following drug concentrations: EMB (2, 4, 8, 16, or 32 μg/ml), INH (0.025, 0.05, 0.1, 0.2, 0.625, 1.25, 2.5, 5 μg/ml), MXF (0.05, 0.1, 0.2, 0.4, 0.5, 1, 2, or 4 μg/ml), and RIF (0.0125, 0.025, 0.05, 0.1, 1, 10, or 100 μg/ml). Antibiotic-free 7H10 agar plates spotted with the primary inoculum or a 1:100 dilution were used as controls. The plates were incubated at 37°C for 2-3 weeks. The first antibiotic concentration that inhibited growth compared to growth of the 1:100 dilution defined the MIC. All MICs were determined in triplicate. Because each value within a triplicate MIC test was almost always identical to the other values within the same triplicate set, a single MIC value is shown without standard deviation for each test. Susceptibility of H37Rv strains to RIF, INH and MXF in 7H9 containing 10% OADC and 0.2% glycerol was determined by the microdilution method, as described previously(55, 56). Briefly, the first column of wells of a 96-well plate (Costar) was filled with 200μl of 7H9 containing the drug at maximum concentration to be tested. The remaining wells were filled with 100μl 7H9 medium. This was used to perform 2-fold serial dilution of the drugs, RIF 0.82-->0.003μg/ml, INH 0.34-->0.0027μg/ml and MXF 1-->0.008μg/ml. Each well was inoculated with approximately 5×10^5^ CFU of M. tuberculosis. Plates were sealed with a Breathe-Easy membrane (Sigma) and incubated at 37°C and shaking for 10 days. The percentage of growth inhibition in each well was determined by measuring optical density at 600nm (OD_600_) in a Cytation 3 Imaging plate Reader (BioTek Instruments) and comparing it to a well with no drug. The susceptibility tests were done in triplicate and statistical significance was analyzed using a two-way ANOVA. Susceptibility of *M. bovis* BCG strains to POA in 7H9 broth at near-neutral pH 6.5 (with 0.5% (vol / vol) or without glycerol) was determined as described previously(53). POA MIC_50_ values reported here are those concentrations of drug that inhibit 50% of growth as compared to a drug-free control after 5 days of incubation. The wild-type strain was included as a control in all experiments. All MICs were performed in three technical and biological replicates.

### Reversion of the small colony phenotype to wild-type colony phenotype

Three independent colonies from small colony phenotype cultures were randomly picked and grown in 7H9 liquid medium at 37°C to late phase (OD_580_ =1.6), which was considered passage 0. Each culture was plated on 7H10 agar medium and the frequency of large colonies was determined. The cultures were then diluted to 8.69 × 10^7^ ± 2.20 × 10^7^ CFU per starting inoculum, and flasks of 12ml 7H9 liquid medium were inoculated. The culture flasks were incubated at 37°C for 1 week. Serial dilutions of the cultures were plated on 7H10 agar medium to determine the CFU count and to examine the emergence of large-colony phenotypes. The culture passages were repeated up to 12 times.

### Growth and colony morphology of POA-resistant *M. bovis* BCG

In order to determine if the mutants had any specific growth-related phenotypes, mid-log cultures of *M. bovis* BCG were pelleted at 3200 rpm for 10 minutes and re-suspended in 7H9 broth (with 0.5% (vol / vol) or without glycerol) by adjusting to a fixed starting inoculum (OD_600_ = 0.03) in T25 culture flasks (SPL Life Sciences) and growth was monitored by measuring OD _600_ at each time point (Ultrospec 10 cell density meter, Amersham Biosciences). Pictures of *M. bovis* BCG colonies were taken under the fixed magnification of a stereomicroscope (Nikon SMZ-Z45T) by a camera (Nikon DS-Fi2) mounted to its eyepiece.

### DNA isolation, PCR, and bidirectional DNA sequencing

Genomic DNA was extracted using CTAB protocol with slight modification as described previously(57, 58). To amplify DNA fragments for DNA sequencing, PCR was performed using a mix containing 1 ng of genomic DNA, 5 pmol of each primer, 200 μM dNTPs, 1x PCR buffer, and 1 U of high-fidelity *pfx Taq* polymerase (Invitrogen) or Phusion DNA polymerase (Thermo Scientific, as per manufacturer’s protocol) per 50μl reaction. All PCR products were examined on an ethidium bromide-stained agarose gel and purified using a gel extraction kit (Qiagen). Direct Sanger sequencing of PCR products was performed with a BigDye Terminator kit and analyzed with an ABI3100 Genetic Analyzer (Applied Biosystems). Sequences for the primers used in this study are available upon request.

### Whole-genome sequencing and mutation detection

Total genomic DNA was extracted as described above and purified by using MagAttract HMW DNA kit (Qiagen). The DNA of TDR193-SC, TDR193-LC, TDR193-SC-RevLC1, TDR193-SC-RevLC9, C71-SC, C71-LC, C71-SC-RevLC, HN878-XM4-LC1, HN878-XM4-LC2, HN878-XM4-LC3, HN878-XM4-SC4, HN878-XM4-SC5, HN878-XM4-SC6, and HN878-SC strains were submitted to The Genomics Center, Rutgers University, Newark, New Jersey, USA (http://research.njms.rutgers.edu/genomics/). DNA libraries were constructed using the Nextera XT DNA Library Preparation Kit (Illumina, San Diego, USA) and samples were sequenced on Illumina MiSeq (Illumina Inc.) to produce more than 100x coverage. The quality trimmed paired-end reads were mapped to *M. tuberculosis* CCDC5079 (GenBank ID: CP002884.1) or to *M. tuberculosis* H37Rv strain (GenBank ID: AL123456.3) and the variants (single nucleotide changes and small deletion up to 50 nucleotides) were detected with the Fixed Ploidy Variant Detection tool in CLC Genomics Workbench version 9 (CLC, Bio-QIAGEN, Aarhus, Denmark). Variants falling in PE/PPE family genes were excluded from the analysis.

### RNA extraction, library preparation, and sequencing

*M. tuberculosis* strains were grown to mid-log phase (OD_600_ =0.6-0.7) at 37°C in 7H9 medium with or without 1% glycerol. Total RNA was extracted by using Trizol reagents (Invitrogen) as described previously(59). The extracted RNAs were submitted for transcriptome analysis to The Genomics Center, Rutgers University, Newark, New Jersey, USA (http://research.njms.rutgers.edu/genomics/). The quality of RNA was checked for integrity on an Agilent 2100 Bioanalyzer using RNA pico6000 kit; samples with RNA integrity number >7.0 were used for subsequent processing. Illumina Ribo-Zero rRNA Removal Kit (Bacteria) was used for the removal of ribosomal RNA according to the manufacturer’s protocol. The Illumina compatible RNAseq library was prepared using NEB next ultra RNAseq library preparation kit. The cDNA libraries were purified using AmpureXP beads and quantified on an Agilent 2100 Bioanalyzer and on Qubit 2.0 Fluorometer (Life Technologies, Carlsbad, CA). An equimolar amount of barcoded libraries were pooled and sequenced on Illumina NextSeq platform (Illumina, San Diego, CA) with 1×75 configuration. CLC Genomics Workbench 10.0.1 version (CLC, Bio-QIAGEN, Aarhus, Denmark) was used for RNA-Seq analysis. De-multiplexed FASTQ files from RNA-Seq libraries were imported into the CLC software. Bases with low quality were trimmed using the following setting: quality trim limit = 0.05, ambiguity trim maximum value = 2. The trimmed reads were mapped to reference genome, *M. tuberculosis* H37Rv (GenBank ID: NC_000962.3). The aligned reads were obtained using the following parameters: maximum number of allowed mismatches was 2, minimum length and similarity fraction was set at 0.8, and minimum number of hits per read was 10. For each strain three independent RNA extractions and RNAseq analysis were performed; and the statistical analysis of differentially expressed genes compared to the control strain is carried out based on a negative binomial GLM model influenced by the multi-factorial EdgeR method(60), a tool in CLC Genomic Workbench.

We ranked RPKM expression values for each sample, such that the lowest RPKM value was designated 1, the second lowest value was designated 2, etc. We used these ranked values as input into the Gene Set Enrichment Analysis (GSEA) Pipeline (http://software.broadinstitute.org/gsea/index.jsp)(61). Using the GSEA analysis software, we tested for enrichment in gene sets based on regulons (information from TFOE(62)). The genes were ranked ordered by the GSEA software according to their ranked RPKM values using the Student’s t statistic metric assessing difference between conditions compared. Gene sets compromising activated and repressed targets genes were analyzed separately, and sets consisting of greater than 10 genes were included in the analysis, and nominal p-values calculated from the analyses of the activated and repressed genes were combined using Fisher’s method(63). A “regulon activity change score” was calculated by weighting the gene-set-size-normalized enrichment score (NES) values of the activated and repressed gene sets with the relative proportions of activated vs. repressed target genes in the regulon. Regulons were considered enriched if GSEA reported enrichment scores (ES) in opposite directions for the activated vs. repressed target gene sets, and if the overall regulon activity change score had an absolute value greater than 1. Among enriched gene sets, those with higher ES, NES, and leading edge statistics were prioritized (see GSEA(61) for more information about the individual statistics).

### Mouse infection, moxifloxacin treatment, and isolation of *glpK* mutants

Pathogen-free female BALB/c mice aged 8 weeks were purchased from Charles River Laboratories (Wilmington, MA). Mice were group-housed in a biosafety level III animal facility and maintained with sterile bedding, water, and mouse chow. All animal experiments were conducted in compliance with and approved by the Investigational Animal Care and Use Committee of the New Jersey Medical School, Rutgers University. Nine-week-old female BALB/c mice (weight range 18-20 g) were infected with an inoculum of 3×10^6^ CFU/mL *M. tuberculosis* HN878 using a Glas-Col whole body aerosol unit. This resulted in a lung implantation of 1.56 log CFU per mouse 3 h post-infection. Groups of 4-5 mice were sacrificed via cervical dislocation 2 and 4 weeks post-infection. At 4 weeks post-infection, two groups of 5 mice were treated daily with 100 mg/kg moxifloxacin (MXF) for one week. One group was analyzed immediately at the end of the one-week treatment period, and the other group was analyzed two weeks later. Control groups received vehicle only for one week. Whole lungs were homogenized in 8 ml of PBS supplemented with 0.05% Tween 80 and serial dilutions were plated on Middlebrook 7H11 agar supplemented with 10% OADC and 0.5% glycerol to score the bacterial burden. Colonies were counted after 6 weeks of incubation at 37 °C.

### Glycerol quantitation in infected lung tissue

Lung tissues were cryosectioned to 25 μm thick and 3,000,000 μm^2^ of sectioned tissue was collected by laser capture microdissection using a Leica LMD 6500(64). Extraction was performed by adding 50μL of acetonitrile to 3,000,000 μm^2^ of 25 μm thick laser captured tissue. Extracts were sonicated in a water batch for 10minutes and centrifuged at 4000 RPM for 5 minutes. Derivatization of glycerol was performed to improve the HPLC retention of glycerol and to reduce spectral background noise. Glycerol derivatization was performed by combining 15 μL of 1% benzoyl chloride derivatization agent in acetonitrile, 15 μL of glycerol standard in acetonitrile or 15 μL of tissue extract, 15 μL of 1 μg/mL glycerol-d5 in methanol and 15 uL of 100 mM sodium carbonate. An additional sample of 15 μL TAG mixture in acetonitrile was placed in the derivatization reaction conditions to demonstrate that glycerol does not deconjugate from TAG species during the reaction and contribute to glycerol levels. The TAG standard contained a total of 10 different species in equal parts and a total concentration of 500μg/mL TAGs. The reaction mixture was heated to 50°C for 1 h to complete derivatization. 60 μL of 2% Formic acid in water was added to the samples to quench the reaction. 5 μL of the standards and study samples was injection on to the AB Sciex 6500+ Qtrap. Chromatography was performed on an Agilent SB-C8 4.6 × 100 mm 3.5 μm HPLC column using a reverse phase gradient. The mobile phases were 0.1% formic acid in Milli-Q deionized water and 0.1% formic acid in acetonitrile. The monobenyzl derivative MRM transitions were monitored for both glycerol (197.08/105.03) and glyderol-d5 (202.08/105.03) internal standard.

### Statistical analysis

All the data were analyzed using a statistical tool of CLC Genomic Workbench or Student’s *t* test, as appropriate. CLC tool was used to analyze RNA-Seq data and differentially expressed mRNAs were defined using False Discovery Rate (FDR) adjusted *p* value ≤0.05. Microsoft Excel was used to perform Student’s *t* test and a *p* value ≤0.05 was considered significant.

### Data availability

The data that support the findings of this study are available from the corresponding authors upon request. Genome and transcriptome data were deposited in the NCBI BioProject database (ID: PRJNA478476). The BioSample accession numbers for genomes and transcriptomes analyzed in this study are described in *SI Appendix*, Table S3.

## Acknowledgements

Supported by the National Institute of Allergy and Infectious Diseases, award numbers U19AI11276 and R01AI111967, and by the Singapore Ministry of Health’s National Medical Research Council under its TCR Flagship grant NMRC/TCR/011-NUHS/2014 and the Center Grant ‘MINE’ Core #4 BSL-3 NMRC/CG/013/2013 to TD and is part of Singapore Programme of Research Investigating New Approaches to Treatment of Tuberculosis (SPRINT-TB, http://www.sprinttb.org). We thank Dr. Anne Lenaerts for providing Kramnic mouse samples and Matthew Zimmerman and Yan Pan for finalizing the glycerol data. We thank Claudia Setzer for help in the initial phase of the project. We thank Drs. Chris Sassetti and Martin Gengenbacher for their helpful discussions.

## Author contributions

H.S., P.G., V.D., T.D. and D.A. designed the studies. H.S., S.L., C.L., P.G., M.Y., L.L., L.B. and H.HL. conducted the experiments. H.S., S.M., T.G., S.H., M.H., P.S., V.D., D.S., T.D. and D.A. analyzed the results. H.S., P.G., S.M., T.D. and D.A. wrote the first drafts, and all authors contributed to and approved the final draft of the manuscript.

